# On the edge of extinction: Delayed plant genetic response to forest edge dynamics

**DOI:** 10.1101/2025.04.23.649530

**Authors:** Lore Hostens, Christophe Metsu, Kasper Van Acker, Gerrit Peeters, Sander de Beer, Yannick Gansemans, Eugenija Kupcinskiene, Lina Jociene, Edvina Krokaite-Kudakiene, Jolina Paulssen, Leonie Mazalla, Martin Diekmann, Timo Conradi, Sarah Plößner, Balázs Deák, Orsolya Valkó, Réka Fekete, Anna Orczewska, Per-Ola Hedwall, Jörg Brunet, Siyu Huang, Jannis Till Feigs, Tobias Naaf, Monika Wulf, Jan Plue, Jaan Liira, Dieter Deforce, Filip Van Nieuwerburgh, Pieter De Frenne, Filip Vandelook, Koenraad Van Meerbeek, Olivier Honnay, Hanne De Kort

## Abstract

Understanding genetic responses to forest dynamics is essential for predicting the long-term viability of understory plant populations and for developing effective conservation strategies. This study investigates genetic extinction debt and colonization credit in *Circaea lutetiana*, a clonal forest understory species, across its European range. Using pooled genotype-by-sequencing data from 40 forest edge and core populations, we examined to what extent population size, latitude and historical changes in forest configuration predict genetic diversity. Our findings reveal that the historical forest configuration profoundly shapes present-day genetic diversity. Long-established forest edge populations exhibit significantly reduced allelic richness (−9%) compared to core populations, indicating the partial pay-off of a genetic extinction debt. In contrast, populations from recently established forest edges maintain comparable allelic richness to core populations, suggesting delayed population genetic responses to land use changes. Finally, populations established in areas that were afforested during the past 250 years exhibit lower genetic diversity than historical forest core populations, indicating a delay in genetic recovery and thus a potential genetic colonization credit. Our results highlight that *C. lutetiana* populations are not at equilibrium with the current forest configuration, underscoring the role of lagged genetic responses across very long time scales. Connectivity and population size further moderate genetic diversity, with smaller, isolated populations particularly vulnerable to genetic erosion. Given the limited research on delayed evolution in forest understory species, our results improve the understanding of extinction risk dynamics and underscore the need for history-informed restoration efforts.

## Introduction

Forests are dynamic ecosystems, shaped by natural disturbances and human activities (McDowell et al., 2020). Over time, these processes influence the spatial structure of forests, leading to the continuous formation and disappearance of forest edges (Fischer et al., 2021; Meeussen et al., 2020). While some edges emerge gradually through natural succession, others form abruptly due to deforestation or afforestation, all creating dynamic boundary zones between forested and non-forested habitats. Notably, at least 20% of the world’s remaining forests are within 100 m of a forest edge, highlighting the widespread occurrence of edge habitats and their potential ecological and evolutionary consequences for forest species (Haddad et al., 2015).

Forest edges are prone to edge effects, which create suboptimal habitat conditions for many shade-adapted forest understory species (Harper et al., 2005). Edge effects significantly alter microclimatic conditions, i.e., vegetation at the edge experiences increased exposure to light, wind, and temperature fluctuations and reduced humidity (Harper et al., 2005; Meeussen et al., 2021a; Murcia, 1995). These altered abiotic conditions can disrupt species interactions, reshape plant community composition and influence the demographic and genetic makeup of populations (Franklin et al., 2021; Harper et al., 2005; Hofmeister et al., 2019; Honnay et al., 2005). For example, pollinator communities are influenced by edge dynamics, which may impact plant reproduction and genetic diversity. While forest edges often support higher pollinator diversity and activity (Ren et al., 2023), they may also favor generalists over specialists, altering pollination interactions and affecting plant population resilience (Spiesman & Inouye, 2013).

Altered pollinator and seed dispersal dynamics in edges can negatively impact plant populations due to Allee effects, inbreeding and reduced colonization rates, all of which can increase the risk of extinction (Browne & Karubian, 2018; Luque et al., 2016; Schlaepfer et al., 2018). For example, small and isolated populations risk losing genetic diversity over time due to genetic drift, when lost alleles are not replenished through gene flow (Honnay & Jacquemyn, 2007; Leigh et al., 2019; Young et al., 1996). In turn, reduced genetic diversity can diminish a population’s ability to adapt to environmental changes and can compromise short-term fitness through inbreeding depression (Ellstrand et al., 1993; Markert et al., 2010). These successive series of demographic and genetic events may cause an extinction vortex, a self-reinforcing decline in genetic diversity and population size that ultimately accelerates the risk of extinction (Figueiredo et al., 2019).

Overall, forest edge understory plant populations may often face an increased extinction risk, yet the extent to which these populations persist over time remains unclear (Harper et al., 2005; Murcia, 1995). Edge populations of long-lived or clonally reproducing species may persist under prominent edge effects due to a delayed response of the population genetic diversity to habitat changes (Honnay & Jacquemyn, 2007). This phenomenon is referred to as a genetic extinction debt, where population genetic diversity has not yet responded, or is still responding, to habitat changes and is expected to decline in the future (Aguilar et al., 2008; Honnay et al., 2005; Tilman et al., 1994). Several studies have found evidence of a genetic extinction debt in clonally reproducing and long-lived plant species where the population genetic diversity was shown to reflect the past rather than the current landscape configuration (Aavik et al., 2019; Plue et al., 2017; Reinula et al., 2021; Reisch et al., 2017). Factors such as distance to the nearest forest, forest size, and population size may all influence the extinction risk of edge populations (González et al., 2020). First, the presence of nearby habitats may facilitate recolonization and delay extinction, while forest isolation may accelerate it. Second, large forests can harbor stable and highly diverse core populations that have not experienced edge effects for centuries, thus enabling genetic rescue of populations in their edges. Gene flow from these core areas or from nearby forests may help slow the loss of genetic diversity in edge populations, thereby delaying population extinction in forest edges (Whiteley et al., 2015). Third, large population sizes may enhance persistence in edges, whereas smaller ones may increase extinction risk (Honnay et al., 2005). Revealing the importance of all these mediating factors will substantially improve our understanding of plant extinction risk under spatial and historical forest dynamics. Recognizing delayed genetic responses also can provide opportunities to restore species’ historical habitat conditions before the negative effects of habitat fragmentation fully appear (Essl et al., 2015).

In contrast to time-delayed genetic erosion, time-lagged genetic responses may also occur as a delayed genetic recovery rate following afforestation. Although these dynamics involve newly established populations, we use the term recovery to emphasize the potential for genetic restoration at the forest or species level. This concept can be referred to as a genetic colonization credit and remains poorly understood. While colonization credit typically refers to species richness, where additional species are expected to colonize a specific area in the future (Baeten et al., 2010; Jackson & Sax, 2010; Nagelkerke et al., 2002), the same concept can indeed be applied to genetic diversity where an increase in genetic diversity is expected following (re)colonization by additional individuals. Such genetic colonization credits are, for example, still visible following the postglacial recolonization of Europe by plant species (Eckert et al., 2008; Hewitt, 1996; Pfeifer et al., 2009). Analogously to the rate of genetic diversity loss, the genetic recovery rate can be expected to depend on the dispersal ability of the plant species, the mobility of its pollinators, landscape connectivity and the number and proximity of source populations (Aavik & Helm, 2018). Whereas some studies have shown that the genetic recovery rate of newly established populations in long-lived plant species can be relatively high under adequate gene flow (Helsen et al., 2013; Jacquemyn et al., 2004), others show that limited gene flow can lead to a rapid loss of genetic variation in founder populations (Jacquemyn et al., 2009).

Here we investigate genetic diversity patterns across European *Circaea lutetiana* L. populations, a clonally reproducing forest understory species occurring in mixed and temperate forests, in relation to historical forest edge changes (up to 250 years ago). To this end, we sampled 40 *C. lutetiana* populations across Europe in both forest cores and edges and employed a pooled genotype-by-sequencing (pool-seq GBS) strategy to obtain allele frequency and heterozygosity data. Mixed models were used to unravel how past and extant forest edge configuration, population size, position within the forest and latitude affect current genetic diversity. We hypothesized that (i) populations in recently created forest edges (i.e. <250 years ago) are not characterized by reduced genetic diversity compared to long-established core populations due to a delayed genetic response, indicating a possible extinction debt, (ii) long-established edge populations exhibit reduced genetic diversity, suggesting that the extinction debt has been at least partially paid off and (iii) newly established populations in areas afforested less than 250 years ago exhibit reduced genetic diversity compared to long-established core populations, indicating a lag in genetic recovery and suggesting the presence of a genetic colonization credit.

## Materials and methods

### Study species

*Circaea lutetiana* (Onagraceae) is a pseudo-annual, clonally reproducing, diploid forest understory herb that occurs in well-drained, deep, loamy soils in deciduous forests but can also be found in shaded hedgerows and riparian zones (Boufford, 1987; Raven, 1963). It is adapted to shade and high air humidity, has a high specific leaf area and is classified as a specialist forest species, indicating that edge habitat might be less suitable (Heinken et al., 2022). Its native range extends across temperate Europe (except N-Europe), North Africa and West Asia (Weeda et al., 1984). Unlike other clonal species, where new ramets emerge within the same growing season, pseudo-annuals produce rhizomes that remain dormant until the next growing season (Jerling, 1988; Verburg et al., 1998). The end part of the rhizome forms a hibernacle towards the end of the growing season, which serves as the primary overwintering organ (Jerling, 1988; Verburg et al., 1996). As autumn progresses, the parent shoot senesces along with the connections between rhizome segments. The next spring, new ramets emerge exclusively from hibernacles that developed during the previous year. Flowers are pollinated by small flies (Diptera), hoverflies (Syrphidae) and to a lesser extent bees of the family Halictidae (Boufford, 1987; Raven 1963). Self-pollination is exceptional (Boufford, 1987). The bristle seeds are dispersed through epizoochory (Lososová et al., 2023).

### Sample collection and demographic variable(s)

We collected leaf material from 40 populations of *C. lutetiana* in the core and at the edge of forest patches of varying sizes (1 ha to 37,608 ha) across 12 regions within the species’ European distribution range (Fig. 1A). The populations of *C. lutetiana* were sampled following a nested design: in each of 12 regions, one or two forests were sampled and within every forest, all populations were sampled with a maximum of five populations per forest. Populations were considered distinct if they were at least 80 m apart and were delineated by taking five to ten coordinates around the populations. Within each population, leaves from a maximum of 300 plants (one leaf per plant) were sampled and individuals were counted (up to 300) or estimated in the field. Population sizes ranged between 48 and 10,000 plants. After collecting, the leaves were dried with silica gel and stored at room temperature.

**Figure 1:**
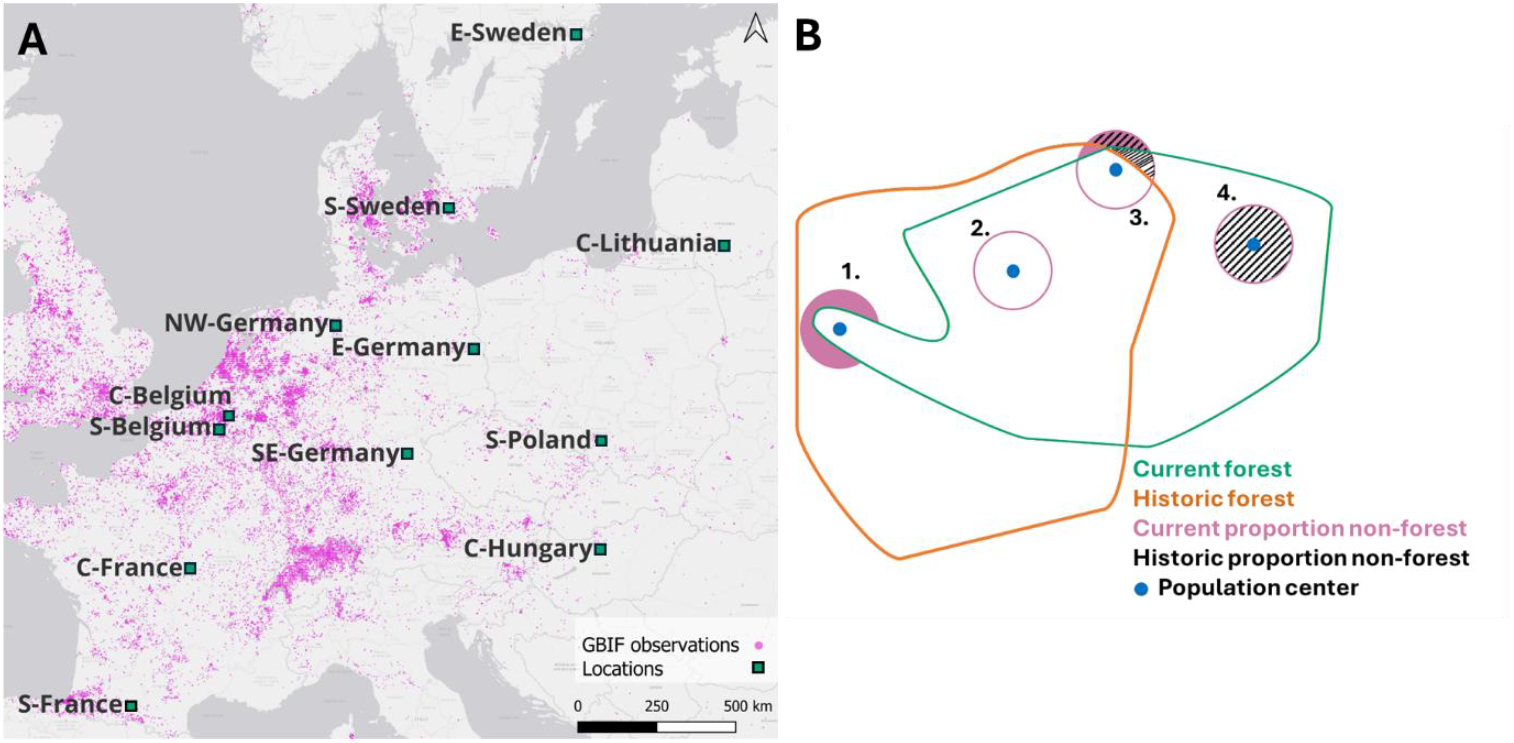
(A) Map of sampling locations (green square) across C. lutetiana’s European range. The pink dots show the species’ range based on the Global Biodiversity Information Facility (GBIF) observations (GBIF.org, 2025). Abbreviations: N = North, E = East, S = South, W = West, NW = North-West, SE = South-East, C = Central. (B) Schematic representation of how the forest dynamics was classified. This figure shows the four categories mentioned in detail in the text below. (1) Recent edge, (2) long-established core, (3) long-established edge and (4) newly established population.

A leaf punch (6 mm) was used to obtain an approximately equal amount of leaf tissue for every individual. This material was pooled within one population and genomic DNA was extracted using a Plant DNA Extraction Kit (Norgen Biotek, 2015). Duplicates of five populations were created for quality control.

### Sequencing, SNP calling and genetic diversity profiling

Double digest (PstI-MseI) genotype-by-sequencing (GBS) using Illumina sequencing was performed by LGC Genomics GmbH to generate libraries for the 40 pooled samples. For all libraries, each sequencing lane was demultiplexed using the Illumina bcl2fastq v2.20 software (Illumina, 2013), allowing one or two mismatches or missing data points in the barcode read. Reads with 5’ ends that did not match the restriction enzyme site were discarded. All reads containing missing data points were removed and reads at 3’ end were trimmed to get a minimum average Phred quality score of 20 over a window of ten bases. Reads with a final length of <20 bases were discarded (Li, 2013).

Quality trimmed reads were aligned against a reference sequence (see data availability) using BWA-MEM v0.7.12. Samples were genotyped and variants were discovered using Freebayes v1.2.0 (Garrison & Marth, 2012). Variants were filtered using the package *vcfR* and *adegenet* in R version 4.2.2 (R Core Team, 2024) using: (i) minimum allele frequency of 5%, (ii) no missing data for maximum accuracy of frequencies and (iii) a minimum total allele count of 200 per SNP. This resulted in a final dataset containing 894 high quality SNPs (Jombart & Ahmed, 2011; Knauss & Grünwald, 2017).

Average allelic richness per population was calculated as the cumulative allele count across all 894 SNPs divided by the total number of SNPs. Estimated within-population heterozygosity was calculated by using the function compute.fstats from the package *poolfstat* (Gautier et al., 2022).

### Spatial variables

To test whether forest patch size, distance to the nearest neighboring forest, exposure to edge effects and latitude affect genetic diversity, we retrieved the (i) **forest area** (1-37,608 ha), (ii) **distance** to the nearest forest patch (5-429 m) as a proxy for connectivity, (iii) the population’s **position** within the forest patch (North, East, South, West, or Central), based on the area where the majority of individuals were concentrated, in order to assess whether edge effects are more pronounced on the southern side due to increased solar radiation (Matlack, 1993) and (iv) **latitude** as a proxy for post-glacial colonization credits. The forest area and distance to the nearest forest patch were calculated using CORINE land cover 2018 (EEA, 2019) and Open Street Map in QGIS version 3.28.2 (QGIS.org, 2024).

To assess how historical changes in forest edge configuration influence the genetic diversity of forest understory populations and to evaluate the presence of genetic extinction debts and colonization credits, we created a categorical variable termed “forest dynamics”. This variable accounts for correlated historical and current landscape configuration. We classified populations into four categories, including “long-established core” (both historically and currently a forest core population), “recent edge” (a population historically in the forest core but currently in the edge), “long-established edge” (both historically and currently a population at the forest edge), or “newly established population” (a new population in an area that was historically not forested) (Fig. 1B). To define the edge, we first generated a 200 m buffer around the population center and calculated the proportion of current and historic non-forest habitat within this buffer. Populations were then classified as recent, newly established or long-established edge or core, based on non-forest habitat thresholds of 0%, 10%, 20%, 30% and 50%. We then selected the most appropriate threshold based on model fit (see below). The historical maps used to calculate the historical non-forest habitat date between 1763 (Central-Hungary) and 1914 (Central-Lithuania). The “forest dynamics” predictor then represents the change in forest cover around the population between 1763 and 2024 (Fig. 1B). Information on the historical maps can be found in Appendix S1.

### Statistics

Allelic richness and heterozygosity were modeled against “forest dynamics” and the above-mentioned additional predictors using generalized linear mixed models (GLMM) implemented in the *lme4* package (Bates et al., 2014) in R (version 4.4.1; R Core Team, 2024) to reveal how much of the variation genetic in genetic diversity can be explained by the spatial variables and population size. Allelic richness was modeled against forest dynamics, population size (ln transformed), latitude, distance to the nearest forest patch (ln) and position using a GLMM with a Gaussian distribution. Forest-ID was used as a random factor to account for random genetic variation associated with the different sampling locations. We ran this model for every forest dynamics threshold (0%, 10%, 20%, 30% and 50% non-forest cover) since it is unknown to which percentage the impact of edge-effects manifest and selected the final model (20% threshold) based on the Akaike Information Criterion (AIC) and marginal R^2^ (Barton, 2024). Because position decreased model fit, we omitted this variable from the models. Multicollinearity and distributional assumptions were not violated according to a model check implemented in the *car* package (Appendix S2) (Fox & Weisberg; 2018). Dunnett’s test from the package *DescTools* was used to compare each forest dynamics group to the long-established core population (Signorell, 2024).

Heterozygosity was modeled against the same predictors using a GLMM with beta distribution in the *glmmTMB* package (Brooks et al., 2017). Assumptions regarding multicollinearity, distribution of the residuals and overdispersion were also checked and not violated using the packages *performance* and *DHARMa* (see Appendix S2 for the values) (Hartig, 2022; Lüdecke et al., 2021). Dunett’s test was also used to compare each forest dynamics group to the long-established core population (Signorell, 2024). Analogously to the allelic richness model, the forest dynamics threshold of 20% without position rendered the best model fit.

## Results

Allelic richness was on average 1.75 (standard deviation (SD) = ±0.12) and ranged from 1.54 (C-Lithuania) to 1.99 (C-Hungary). Heterozygosity was on average 0.184 (SD = ±0.079) and ranged from 0.083 (SE-Germany) to 0.317 (C-Hungary). Averages and SD’s for both allelic richness and heterozygosity in every forest dynamics group can be found in Table 1. The per-forest difference in allelic richness between long-established core populations and all other forest dynamics groups was 3.1% (SD ± 6.1) for recent populations, 6.4% (SD ± 8) for long-established edge populations and 6.7% (SD ± 6) for newly established populations. For heterozygosity these were 27% (SD ± 46), 2.9% (SD ± 36) and 34% (SD ± 36), respectively. We also found no clear patterns in relatedness between populations found in the same forest (Appendix S3), suggesting that populations within forests are genetically isolated.

**Table 1:** Mean allelic richness (AR) and heterozygosity (hetero) of C. lutetiana with standard deviations (SD) for different forest dynamics categories. Abbreviations: LE = long-established, NE : newly established.

| Forest dynamics | Mean AR | SD AR | Mean hetero | SD hetero |
| --- | --- | --- | --- | --- |
| LE core | 1.83 | 0.1 | 0.23 | 0.07 |
| LE edge | 1.71 | 0.11 | 0.16 | 0.08 |
| Recent edge | 1.77 | 0.06 | 0.22 | 0.04 |
| NE | 1.7 | 0.13 | 0.15 | 0.08 |

The GLMM, which also accounts for demographic and landscape variables, explained 39% (predictors: R^2^_m_) and 66% (full model including random factor: R^2^_c_) of the variance in allelic richness and revealed significant effects of forest dynamics and population size (Appendix S4). Post-hoc analysis using Dunnett’s test indicated that there was no significant difference in allelic richness between long-established core populations and those that recently became edge populations (mean difference = −0.056, SE = 0.073, t(29) = −0.737, p = 0.78) (Fig. 2A). This is in line with our first hypothesis that recent forest edge populations are not characterized by reduced allelic richness. Long-established edge and newly established populations exhibited reduced allelic richness, supporting our second and third hypothesis. Long-established core populations had significantly higher allelic richness compared to long-established edge populations (mean difference = −0.12, standard error (SE) = 0.035, t(26) = −2.54, p = 0.032) (Fig. 2A) and newly established populations (mean difference = −0.13, SE = 0.042, t(32) = −2.57, p = 0.019) (Fig. 2A). Standardized coefficients for all predictors are reported in Appendix S4.

**Figure 2:**
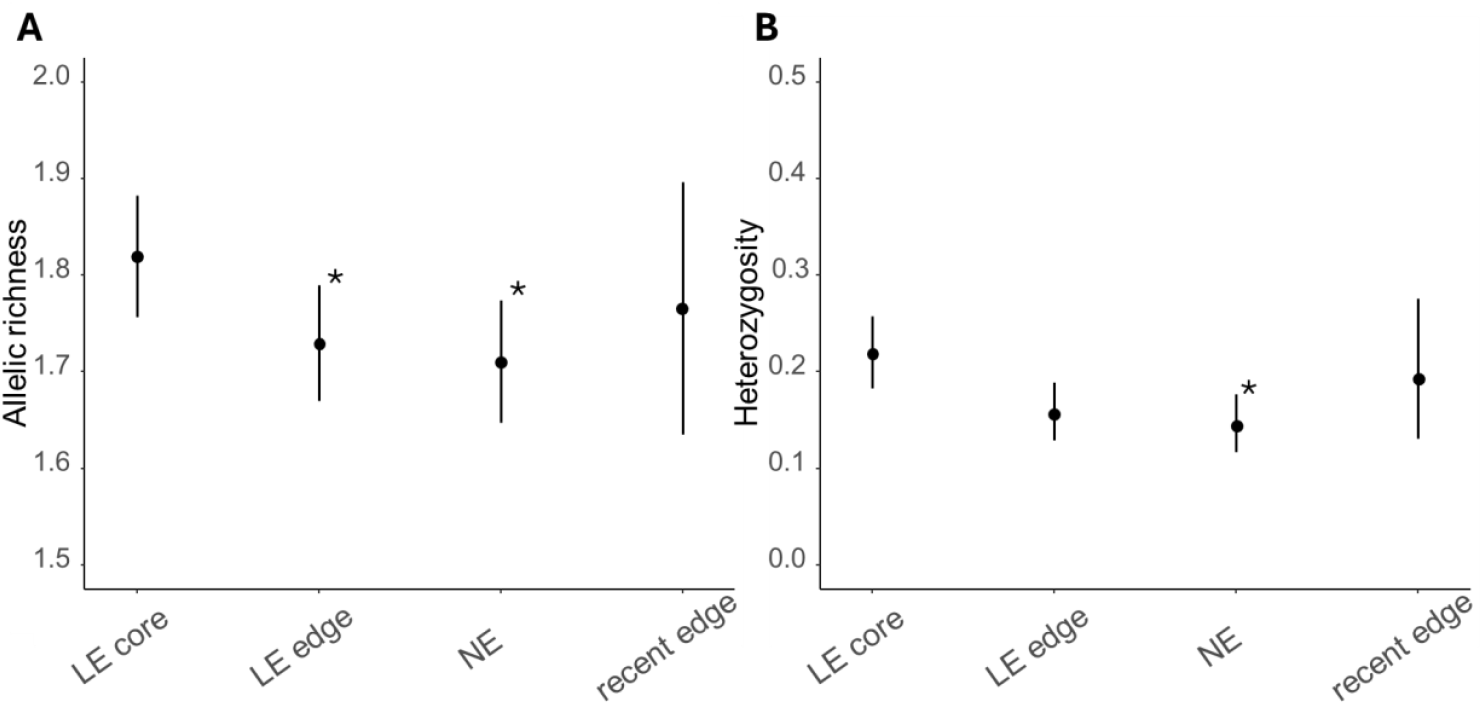
Prediction of the (A) allelic richness and (B) heterozygosity of C. lutetiana in relation to the forest dynamics based on the GLMMs. Black dots show the average genetic diversity per individual factor, whereas error bars show the standard deviation. Abbreviations: LE = long-established, NE = newly established. Asterisks highlight the significant difference in genetic diversity compared to the baseline (long-established core) (*p ≤ 0.05).

Moreover, the allelic richness of *C. lutetiana* populations decreased significantly with decreasing population size (β = 0.30, 95% confidence interval (CI) [0.03, 0.57], t(32) = 2.31, p = 0.028) (Fig. 3A), suggesting that smaller populations hold less genetic diversity. The allelic richness also decreased marginally with increasing latitude (β = −0.33, 95% CI [−0.7, 0.05], t(18) = −1.79, p = 0.091) (Fig. 3C), possibly reflecting post-glacial colonization patterns. None of the other variables significantly influenced the allelic richness (Fig. 3E; Appendix S4).

**Figure 3:**
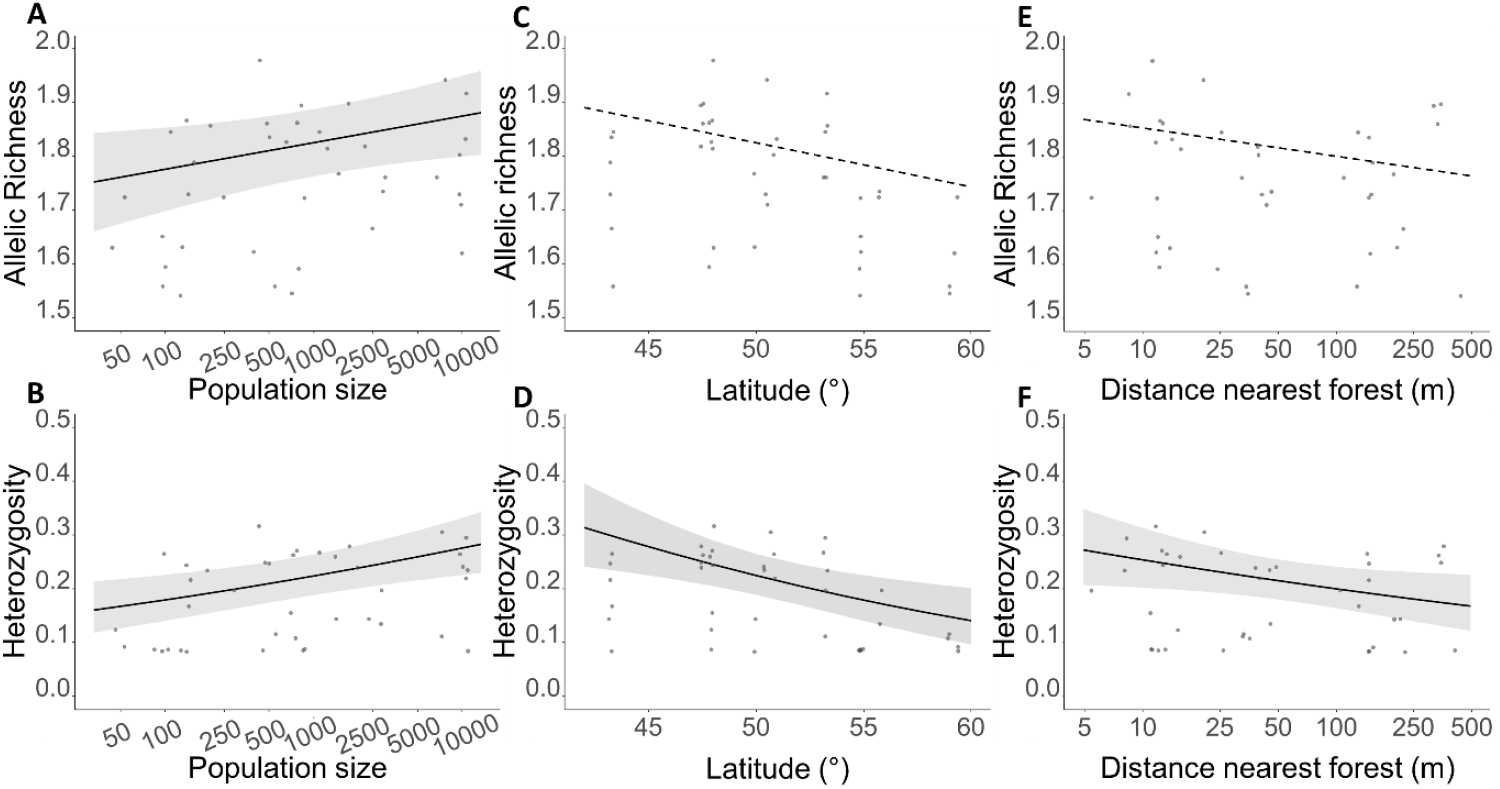
Marginal effects plots of the allelic richness and heterozygosity of C. lutetiana as a function of (A/B) the population size, (C/D) latitude and (E/F) the distance to the nearest forest (m). The grey area represents the 95% confidence interval (CI), while the grey dots indicate the individual raw data points. Dashed trend line and no CI portray non-significance.

Estimated heterozygosity rendered similar patterns. Specifically, the GLMM explained 60% (R^2^_m_) and 84% (R^2^_c_) of the variance in heterozygosity and revealed significant effects of forest dynamics, population size, latitude and distance to the nearest forest patch. Post-hoc analysis using Dunnett’s test indicated that there was no significant difference in heterozygosity between long-established core populations and recent edge populations (mean difference = −0.007, SE = 0.27, z = −0.560, p = 0.99) (Fig. 2B), again in line with our first hypothesis of a delayed genetic response to habitat changes. Although long-established forest edge populations had a marginally lower heterozygosity in contrast to long established core populations, this difference was not statistically significant (mean difference = −0.062, SE = 0.13, z = −3.05, p = 0.11) (Fig. 2B). However, long-established core populations had a significantly higher heterozygosity compared to newly established populations (mean difference = −0.076, SE = 0.16, z = −3.14, p = 0.019) (Fig. 2B), further supporting our third hypothesis that a genetic colonization credit exists in newly formed forest patches. Standardized coefficients for all predictors are reported in Appendix S4.

Furthermore, the heterozygosity of *C. lutetiana* populations decreased significantly with decreasing population size (β = 0.20, 95% CI [0.07, 0.33], z = 3.12, p = 0.002) (Fig. 3B), with increasing latitude (β = −0.27, 95% CI [−0.45, −0.10], z = −3.06, p = 0.002) (Fig. 3D) and with increasing distance to the nearest forest patch (β = −0.17, 95% CI [−0.34, −0.01], p = 0.038) (Fig. 3E). The latter suggests some gene flow between forests. Forest area did not significantly influence the heterozygosity (Appendix S4).

## Discussion

This study investigated the extent of delayed genetic responses of populations in forest edges to changes in forest cover throughout the European range of *C. lutetiana*. Using pool-seq GBS data from 40 populations sampled in forest cores and edges, we assessed how population size, distance to nearby forests, latitude and changes in landscape configuration affect genetic diversity. Evidence for delays in the genetic response of plant populations to land use changes is still scarce and the influence of historical habitat configuration is often overlooked (Aavik et al., 2019; Plue et al., 2017; Reinula et al., 2021; Reisch et al., 2017). Our results show that current genetic diversity is predominantly shaped by landscape-scale forest cover dynamics, suggesting the presence of genetic extinction debts and colonization credits. Our findings indicate that: (i) recent edge populations are not characterized by reduced genetic diversity compared to long-established core populations due to a delayed genetic response, indicative of a genetic extinction debt (ii) long-established edge populations exhibit significantly lower genetic diversity, suggesting a partial pay-off of the genetic extinction debt, and (iii) newly established populations in recently afforested areas possess significantly lower genetic diversity than long-established core populations, indicating a delay in genetic recovery and thus the presence of a possible genetic colonization credit. In Europe, broadleaved forests cover approximately 118 million hectares, with a total edge perimeter of about 9.4 million km (Meeussen et al., 2021b). This emphasizes widespread occurrence of edge effects and their delayed genetic consequences on forest understory species.

### Delayed genetic responses in forest edges

Previous studies have shown that plant populations can retain high genetic diversity after habitat loss and fragmentation (Hahn et al., 2013; Honnay et al. 2006; Mhemmed et al., 2008). Clonally reproducing species, such as *C. lutetiana*, may exhibit delayed genetic responses to habitat alterations (Honnay et al., 2005). The confirmation of our first and second hypothesis support this idea: we observed (i) no significant difference in genetic diversity between long-established core populations and recent edge populations (i.e. former core populations that became edge populations in the last 110-250 years) and (ii) reduced genetic diversity in long-established edge populations, particularly regarding allelic richness. This suggests that long-established edge populations may have already (partially) paid the extinction debt whereas recent edge populations carry an extinction debt that can be expected to become apparent over time. It may therefore take hundreds of years for *C. lutetiana* populations to pay off an extinction debt, irrespective of the size of the forest. Our findings reinforce the notion that genetic erosion occurs over long periods of time, underscoring the necessity of historical context in genetic studies (Epps & Keyghobadi, 2015).

Other studies also found evidence that historical habitat configuration plays an important role in the current genetic makeup of plant populations (Aavik et al., 2019; Münzbergová et al., 2013; Reinula et al., 2021; Reisch et al., 2017). Reinula et al. (2021) reported that the genetic diversity of *Primula veris* L. was significantly shaped by historical grassland edge density and the proportion of historical grasslands. Similarly, Aavik et al. (2019) found that in *Trifolium montanum* L., heterozygosity was influenced by historical grassland area. These findings highlight the potential for delayed genetic responses to changes in the landscape, a pattern also evident in our study.

### Colonization credit

Newly established populations in areas that were afforested in the past 250 years exhibited significantly lower genetic diversity than long-established core populations, which is not in line with some earlier reports on the more rapid genetic recovery of newly founded populations of long-lived or clonally reproducing herbs (Helsen et al., 2013; Huang et al., 2024; Jacquemyn et al., 2004). The significantly lower genetic diversity in populations of *C. lutetiana* that established in the recent past indicates a delayed genetic response to afforestation, reflecting a genetic colonization credit. This pattern is likely driven by limited seed dispersal, genetic drift and restricted pollen flow from nearby populations. Natural recovery without active restoration measures (i.e., seeding) depends on species’ dispersal capacity, landscape connectivity and the availability of nearby source populations (Aavik & Helm, 2018).

Notably, Reynolds et al. (2013) found that the natural recovery of marine *Zostera marina* L. populations, a clonally reproducing plant species, would take between 128-145 years to attain the same level of genetic diversity observed in populations established through seeding in only 10 years. This suggests that natural recovery occurs at a very slow rate in dispersal-limited landscapes, which supports our own findings. Intense genetic colonization credits have also been associated with post-glacial colonization (e.g. Widmer & Lexer, 2001). Correspondingly, our results show a decrease in genetic diversity with increasing latitude, suggesting a post-glacial genetic colonization credit. As populations migrated northward from southern glacial refugia following the last Ice Age, they experienced sequential bottlenecks, resulting in a gradual reduction of genetic variation over successive generations (Hewitt, 1996; Pfeifer et al., 2009; Widmer & Lexer, 2001).

Pronounced genetic delays, particularly where historical maps date back hundreds of years, suggest that clonal species can bridge very long periods of habitat deterioration and that genetic recovery is an extremely slow process. Such delayed genetic responses of forest edge populations highlight the vulnerability of many forest populations across Europe to the risks of habitat changes, particularly through edge effects.

### Effect of connectivity and population size on genetic diversity

Genetic diversity decreased with increasing isolation from other forest patches. Given that *C. lutetiana* is insect-pollinated, this suggests that forest fragmentation may impede pollen-mediated gene flow between populations, consequently accelerating the loss of genetic diversity (Aguilar et al., 2019; Auffret et al., 2017; Honnay et al., 2005). Additionally, although *C. lutetiana* seeds may be dispersed over long distances through epizoochory, the low connectivity of the landscape can impede the movement of organisms, thereby reducing seed dispersal and gene flow between fragments (Mony et al., 2018; Taylor et al., 1993). This is in line with a recent study demonstrating global reductions in plant dispersal through defaunation (Fricke et al. 2021) and highlights the importance of maintaining connectivity between forest patches to mitigate genetic erosion.

Although clonal species can reach high population densities of low genetic diversity due to their clonality, drift and breeding between related individuals (Ellstrand & Elam, 1993; Honnay & Jacquemyn, 2007; Jacquemyn et al., 2006), we demonstrate a significant positive correlation between census population size and genetic diversity across *C. lutetiana* populations at a range-wide scale. This correlation is expected where widespread forest fragmentation causes extinction vortices characterized by demographic and genetic delay, and corresponds to the rationale that clonal species can sustain high population densities in suboptimal conditions for long time periods (Jacquemyn et al., 2006). However, we found no significant relationship between population size and forest dynamics as revealed by additional modeling (Appendix S4)(Brooks et al., 2017), suggesting that demographic patterns do not reflect extinction debt in clonal species. Instead, genetic analyses remain imperative for capturing the long-term consequences of habitat changes on population viability and for providing deeper insights into extinction risk dynamics.

There are a few considerations to note in our study. Due to high collinearity between historical and current landscape configuration, an unavoidable issue, we categorized landscape changes, which reduces the amount of genetic variation that we can explain by forest dynamics. We nevertheless found significant historical edge effects that likely underestimate the actual extinction debt. We further assumed that the age of a population is correlated with the age of the forests, which we could roughly infer from historical maps, an assumption that may have caused additional noise to our results. Lastly, pool-seq GBS allowed us to efficiently analyze a large number of samples, providing valuable insights into the genetic diversity across populations. However, this method may overlook rare alleles, subsequently underestimating the loss of alleles in edge populations. Overall, we are confident that our findings are conservative and underpin substantial genetic extinction debts and colonization credits in a currently common forest understory species.

### Potential conservation implications

The use of contemporary genetic data in conservation assumes that such data accurately reflects the current status of a population. While this is generally true, the current genetic state may obscure ongoing genetic changes that have yet to manifest due to extinction debts and colonization credits. As our results indicate, genetic parameters can exhibit delayed responses to new environmental conditions or disturbances (Epps & Keyghobadi, 2015; Essl et al., 2015; Watts et al., 2020). Ignoring these genetic delays can lead to inaccurate conservation assessments and inefficient allocation of resources for biodiversity protection.

While *C. lutetiana* is not an obligate old-growth forest understory species, our findings suggest delayed genetic responses across its European range. These results also highlight opportunities for conservation and restoration efforts to mitigate further genetic loss and prevent the full realization of the extinction debt (Essl et al., 2015). First and foremost, efforts should be made to prevent further habitat fragmentation in order to preserve core populations and populations with a colonization credit. To safeguard the genetic diversity of forest understory species, preserving the integrity of well-connected forests with core habitat is crucial. Furthermore, our findings suggest that restoration efforts will be most rapid in recently fragmented forests, where full genetic recovery can be anticipated in the short term. Neither forest size nor connectivity had notable effects on allelic richness, however, suggesting that directly dealing with edge effects is the most effective restoration strategy. We specifically recommend targeted reforestation and strategically reconnecting forest fragments to reduce edge effects while improving pollinator movement between populations. This is supported by Brunet et al. (2011) who found that *C. lutetiana* was most frequent in post-arable forests connected to older forest patches with established source populations. Similarly, forest patch size or age had no significant effect on the species’ occurrence, further emphasizing the importance of proximity to source populations in core habitat for rapid population establishment and expansion (Brunet et al. 2021).

Long-established edge populations already paid off much of their extinction debt, leaving genetically depleted populations with an enhanced risk of extinction. Passive genetic recovery following restoration is a slow process. However, if the goal is to minimize genetic losses, priority should be given to populations inhabiting historical forest edges. This recovery process may be accelerated through active interventions, such as population reinforcement via seeding from nearby populations (Reynolds et al., 2013). To what extent historical forest edges harbor high forest species richness with a substantial extinction risk has yet to be established, but our findings raise concern for obligate forest understory species, which are likely to be more affected by habitat change and its consequences (Honnay et al., 2005).

## Conclusion

This study emphasizes the crucial role of historical forest dynamics in shaping the genetic diversity of forest understory species. Past landscape changes, including habitat fragmentation and afforestation, significantly influence current genetic patterns. Our results suggest that these historical processes are key drivers of genetic extinction debts and colonization credits, which must be considered in restoration efforts.

We have established that historical forest edges are particularly vulnerable to delayed evolution in *Circaea lutetiana* populations across Europe. Long-established edge populations exhibit notably lower genetic diversity than core populations, while recent edge populations do not show this difference, suggesting a possible genetic extinction debt. Recently established populations also show lower genetic diversity compared to long-established core populations, indicating a potential genetic colonization credit. Our findings highlight the influence of landscape-scale forest cover dynamics on genetic diversity and emphasize the importance of preserving large, well-connected forests. Moreover, they underscore the need for landscape-scale restoration of aged forest edges, which face a higher extinction risk due to the partial realization of the extinction debt. Given the limited research on delayed evolution in forest understory species, our results improve the understanding of extinction risk dynamics and highlight the need for cautious history-informed restoration efforts, especially for obligate understory species.

## Data availability

All data and code supporting the findings of this study are available on [Zenodo](https://zenodo.org/records/15130887?preview=1&token=eyJhbGciOiJIUzUxMiJ9.eyJpZCI6ImYzY2NiMjk2LWFkYjgtNDViMy05N2FjLWVmNDAxMGU5M2VhNyIsImRhdGEiOnt9LCJyYW5kb20iOiIwNTkzMzY1MzRlN2RjZWVjMmRmNWMyM2Y2YTlmZmIwZiJ9.3uIIge71g0HRGzBL0Y-Ub8TJze2a9jnSMLA8-S5FwUHBmPjsAhPZup5eCZ2Tnr8z7dCt9hzm0tCN2j3O71kyqA). The raw sequence reads of the populations have been deposited in the NCBI Sequence Read Archive under BioProject accession PRJNA1249044. The reference sequence used to align the reads has been deposited at DDBJ/ENA/GenBank under the accession JAYWLA000000000.

## Acknowledgments

This work was supported by the Research Foundation – Flanders (FWO) by funding the scientific research network FLEUR (W000322N, www.fleur.ugent.be). Anna Orczewska would like to thank to Adrian Król for his help in the fieldwork. Balázs Deák, Réka Fekete, and Orsolya Valkó also express their gratitude to Eszter Korom for her help in the field.

