## Supporting information for "On the edge of extinction: Delayed plant genetic response to forest edge dynamics"

### Appendix S1: Historical maps

Table S1: Location, name, year and resolution of the historical map used to create the forest dynamics categories.

| Location | Name | Scale | Year |
| --- | --- | --- | --- |
| C-Belgium | Cartes de Ferraris | 1 : 11 520 | 1770-1778 |
| C-France | Carte de Cassini | 1 : 86 400 | 1786-1815 |
| C-Hungary | Königreich Ungarn [B IX a 527] - First Military Survey | 1 : 28 800 | 1763-1787 |
| C-Lithuania | Kauno Zapyskis Meta duomenys | 1 : 84 000 | 1910 |
| E-Germany | Schmettausches Kartenwerk | 1 : 50 000 | 1776-1787 |
| E-Sweden | Häradsekonomiska kartan | 1 : 20 000 | 1901-1906 |
| NW-Germany | Kurhannoversche Landesaufnahme des 18. Jahrhunderts | 1 : 25 000 | 1764-1786 |
| S-Belgium | Cartes de Ferraris | 1 : 11 520 | 1770-1778 |
| SE-Germany | Historische Karte von Bayreuth-Birken | 1 : 2500 | 1811-1850 |
| S-France | Carte de Cassini | 1 : 86 400 | 1786-1815 |
| S-Poland | Gleiwitz. Karte des Deutschen Reiches. Reichsamt fur Landesaufnahme | 1 : 100 000 | 1893 |
| S-Sweden | Uppmätt av Fältmätningsbrigaden | 1 : 20 000 | 1815 |

### Appendix S2: Model assumptions

1. GLMM allelic richness

The assumptions of the GLMM model for allelic richness were assessed and not violated. The residuals vs. fitted plot indicated no significant deviations, suggesting that the assumptions of linearity and homoscedasticity were met (Fig. S1A). The Q-Q plot (Fig. S1B) and the Shapiro-Wilk test (W=0.97; p=0.34) showed that the residuals were approximately normally distributed. Variance Inflation Factors (VIF) for the fixed effects were below 5, indicating no multicollinearity (Table S2). Furthermore, no significant outliers or overly influential observations were detected based on the Bonferroni test for outliers.

Table S2: Variance Inflation Factor results for the fixed effects of the GLMM for allelic richness.

|  | **Area forest** | **Forest dynamics** | **Nearest forest** | **Latitude** | **Population size** |
| --- | --- | --- | --- | --- | --- |
| GVIF | 1.55 | 1.68 | 1.27 | 1.41 | 1.12 |
| Df | 1 | 3 | 1 | 1 | 1 |
| GVIF^(1/(2*Df)) | 1.24 | 1.09 | 1.13 | 1.19 | 1.06 |

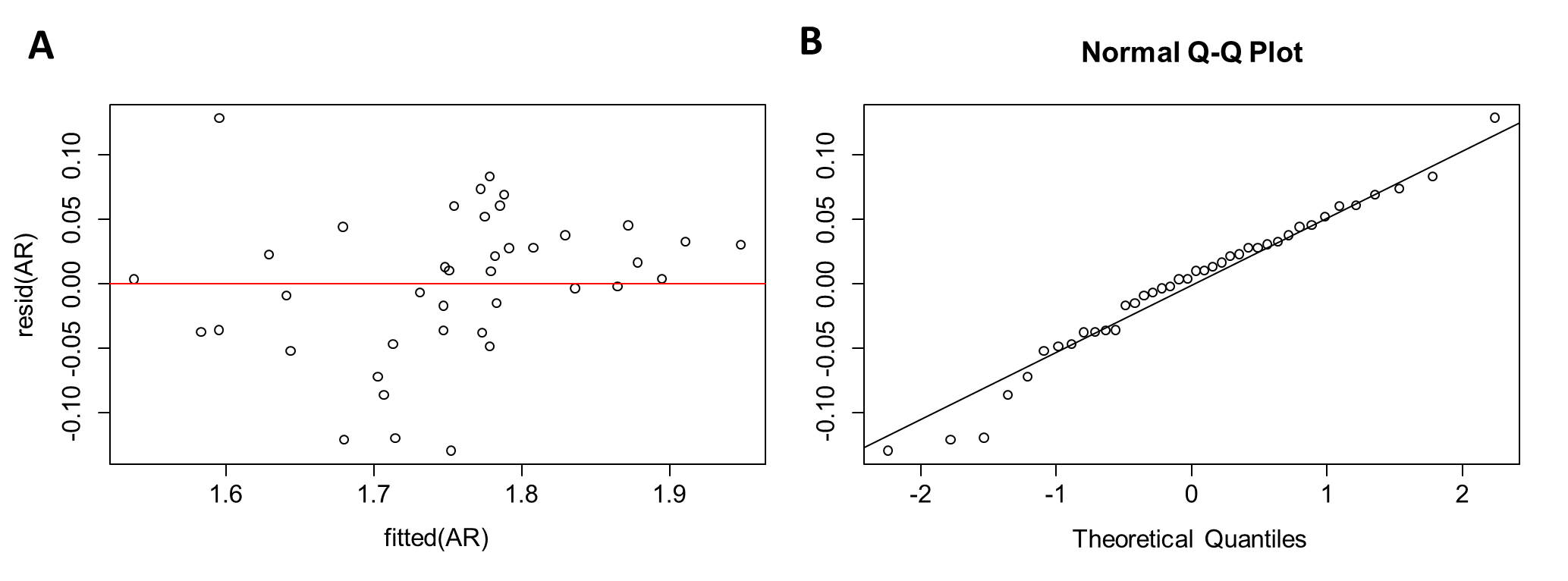

Figure S1: (A) The residuals vs. fitted and (B) Q-Q diagnostics plot of the GLMM model for allelic richness.

2. GLMM heterozygosity

The assumptions of the GLMM model for estimated heterozygosity were assessed and not violated. The dispersion test using simulated residuals from *DHARMa* was non-significant (p=0.85), indicating no overdispersion. The Kolmogorov-Smirnov test for residuals indicated no significant deviation from uniformity (p=0.68) and the residuals vs. predicted plot did not reveal any systematic patterns, suggesting that the model's assumptions were met (Fig. S2). Lastly, multicollinearity among fixed effects was evaluated and all Variance Inflation Factors (VIFs) were below 5 (Table S3), indicating no multicollinearity.

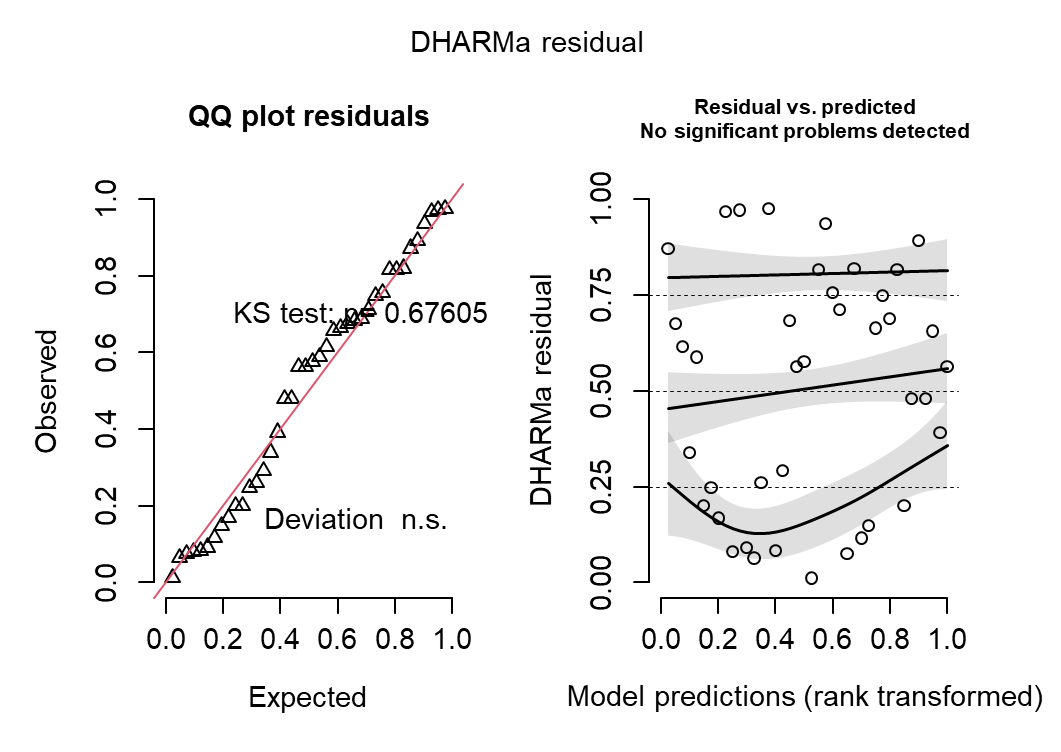

Figure S2: Kolmogorov-Smirnov test for residuals (left) and residuals vs. predicted plot (right) of the GLMM for estimated heterozygosity.

Table S3: Variance Inflation Factor results for the fixed effects of the GLMM for estimated heterozygosity.

| **Term** | **Area forest** | **Forest dynamics** | **Nearest forest** | **Population size** | **Latitude** |
| --- | --- | --- | --- | --- | --- |
| VIF | 1.51 | 1.55 | 1.22 | 1.13 | 1.27 |
| VIF 95% CI | [1.21, 2.24] | [1.23, 2.30] | [1.05, 1.95] | [1.01, 2.16] | [1.08, 1.97] |
| Increased SE | 1.23 | 1.25 | 1.11 | 1.06 | 1.13 |
| Tolerance | 0.66 | 0.64 | 0.82 | 0.89 | 0.78 |
| Tolerance 95% CI | [0.45, 0.83] | [0.44, 0.81] | [0.51, 0.95] | [0.46, 0.99] | [0.51, 0.93] |

3. GLMM population size

The assumptions of the GLMM model for population size were assessed and not violated. Since a negative binomial model was used, overdispersion was explicitly modeled and did not require additional testing. The Kolmogorov-Smirnov test for residuals indicated no significant deviation from uniformity (p=0.23) and the residuals vs. predicted plot did not reveal any systematic patterns, suggesting that the model's assumptions were met (Fig. S3). Lastly, multicollinearity among fixed effects was evaluated and all Variance Inflation Factors (VIFs) were below 5 (Table S4), indicating no multicollinearity.

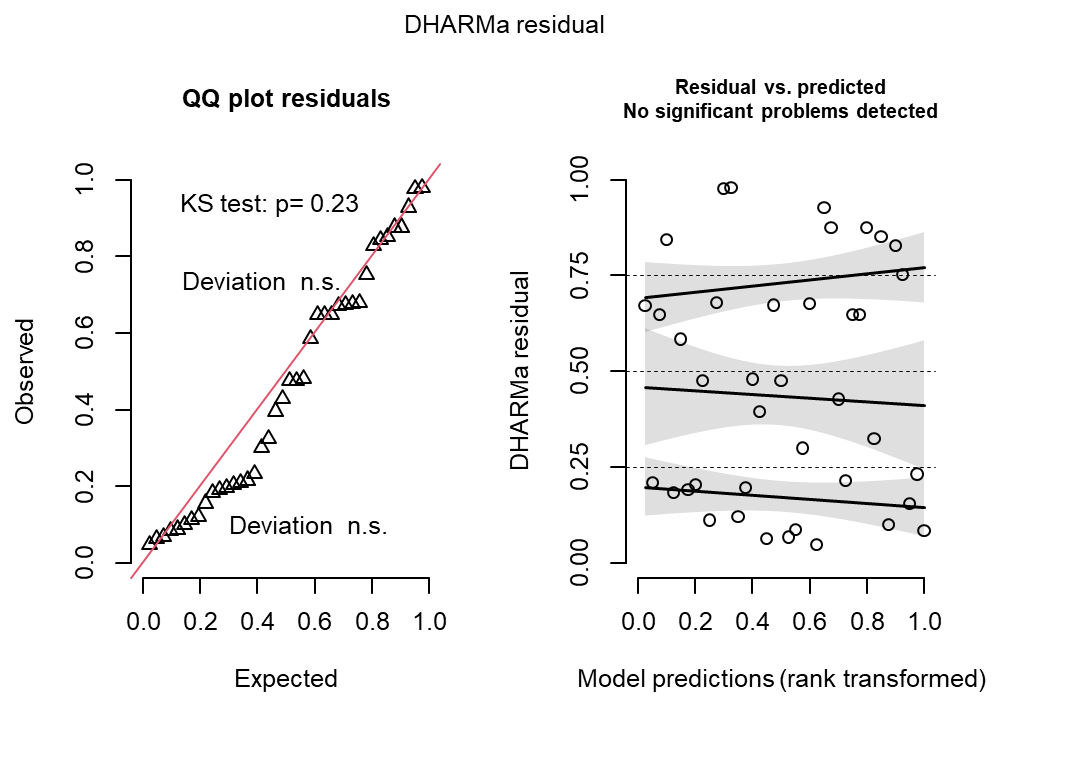

Figure S3: Kolmogorov-Smirnov test for residuals (left) and residuals vs. predicted plot (right) of the GLMM for population size.

Table S4: Variance Inflation Factor results for the fixed effects of the GLMM for population size.

| **Term** | **Area forest** | **Forest dynamics** | **Nearest forest** | **Latitude** |
| --- | --- | --- | --- | --- |
| VIF | 1.23 | 1.15 | 1.34 | 1.15 |
| VIF 95% CI | [1.05, 2.00] | [1.02, 2.10] | [1.11, 2.07] | [1.02, 2.10] |
| Increased SE | 1.11 | 1.07 | 1.16 | 1.07 |
| Tolerance | 0.81 | 0.87 | 0.74 | 0.87 |
| Tolerance 95% CI | [0.50, 0.95] | [0.48, 0.98] | [0.48, 0.90] | [0.48, 0.98] |

### Appendix S3: Relatedness

The PCA based on allele frequencies of all SNPs within the populations shows no clear genetic clustering among populations, as populations from different geographic regions are widely dispersed along PC1. While some populations exhibit slight differentiation, there is substantial genetic overlap between regions (Fig. S4)

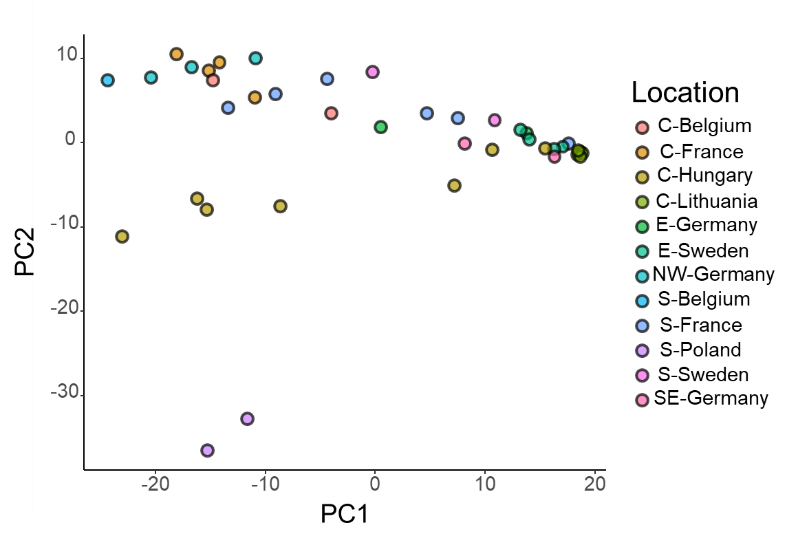

Figure S4: Principal Component Analysis (PCA) of allele frequencies based on pooled-sequencing data across all sampled populations. Each point represents a population, colored by geographic location. Abbreviations: N = North, E = East, S = South, W = West, NW = North-West, SE = South-East, C = Central.

The heatmap of pairwise Fst values among populations within the same forest reveals substantial variation in genetic differentiation (Fig. S5). Populations from the same forest do not always exhibit similar levels of genetic diversity, as indicated by varying Fst values across different population pairs. Notably, some populations show high genetic similarity (low Fst), whereas others exhibit strong genetic differentiation (high Fst), suggesting that even within the same forest there is genetic differentiation.

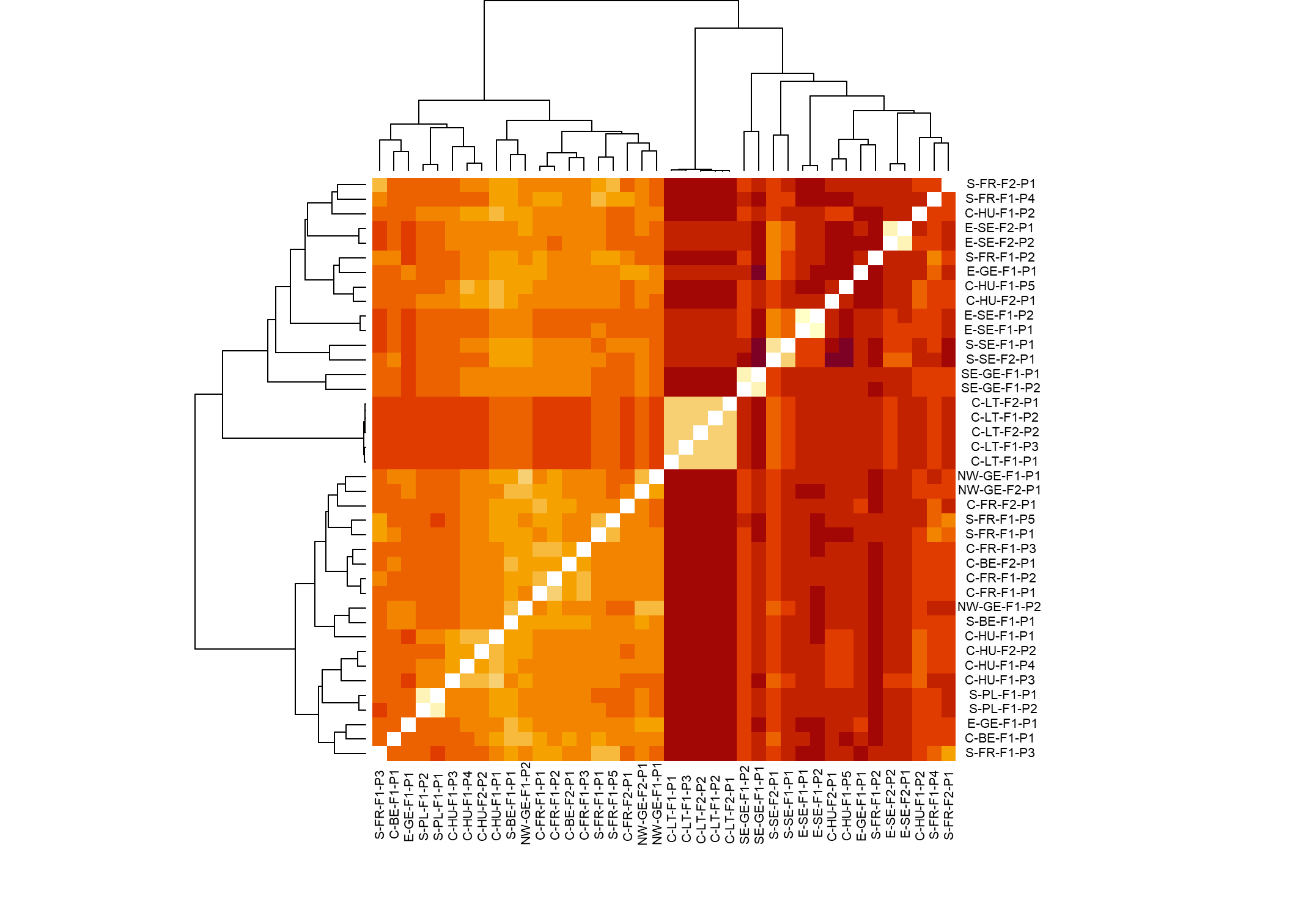

Figure S5: Heatmap showing pairwise Fst values. The lighter the colors indicate higher similarities (lower Fst values). The hierarchical clustering dendrograms (above and to the left) group populations based on genetic similarity, revealing patterns of genetic structure. Abbreviations: N = North, E = East, S = South, W = West, NW = North-West, SE = South-East, C = Central, BE= Belgium, FR = France, GE = Germany, HU = Hungary, LT = Lithuania, PL = Poland, SE = Sweden, F = forest, P = population. Clustering was performed using hierarchical clustering based on pairwise Fst values computed with poolfstat (Gautier et al., 2022).

### Appendix S4: Model outcomes

Table S5: Fixed effect standardized coefficients of the generalized linear mixed models (allelic richness, heterozygosity and population size) (*p ≤ 0.05, **p ≤ 0.01, ***p ≤ 0.001). The factor used for the intercept was long-established core and all other factors were tested against this baseline. Abbreviations: LE = long-establishes, NE = newly established

| **Model** | **Intercept  (LE core)** | **Area** | **LE edge** | **NE** | **Recent edge** | **Latitude** | **Nearest  forest** | **Population size** |
| --- | --- | --- | --- | --- | --- | --- | --- | --- |
| Allelic richness | 0.57*** | 0.08 | -0.76* | -0.92* | -0.45 | -0.33 | -0.25 | 0.30* |
| Hetero-zygosity | -1.28 | -0.06 | -0.40** | -0.50** | -0.15 | -0.27 ** | -0.17* | 0.20** |
| Population size | 7.4*** | -0.28 | -0.47 | 0.47 | 0.31 | 0.13 | -0.41 | / |

Population size is highly variable, particularly in newly established and recent edge populations (Fig. S6A). Distance to the nearest forest is generally small but varies most in newly established populations (Fig. S6B). Forest area follows a decreasing trend from long-established core to recent edge populations, with newly established populations showing high variability (Fig. S6C). This highlights differences in habitat characteristics among population types.

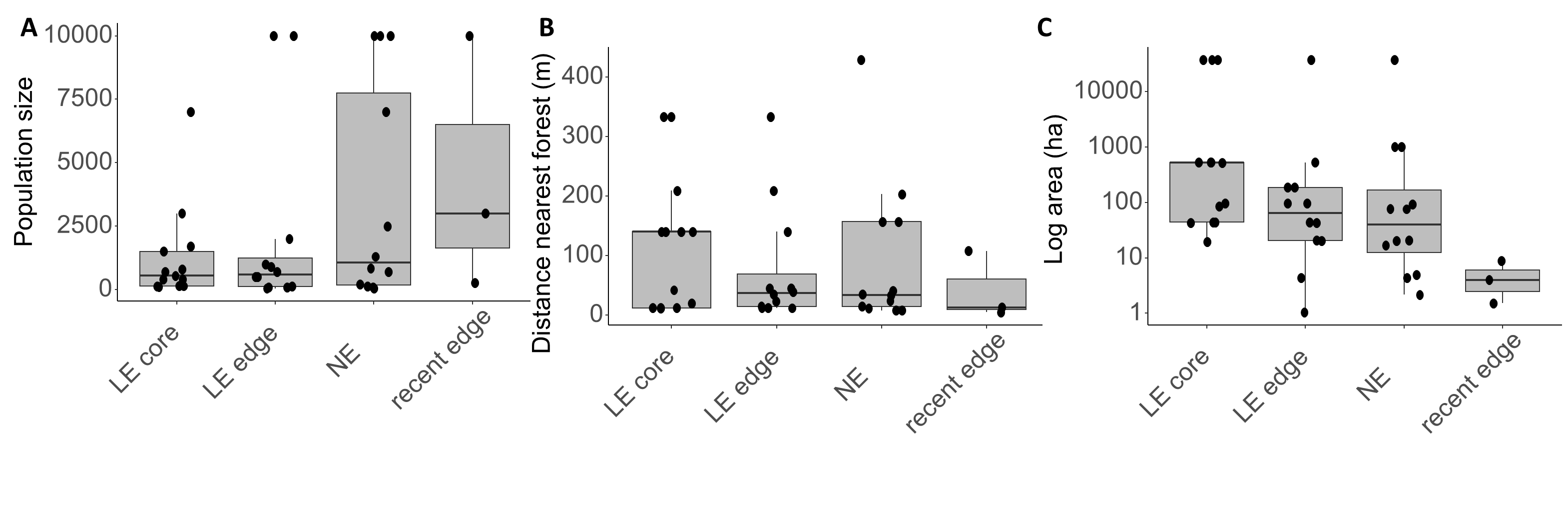

Figure S6: Boxplots showing the distribution of (A) population size, (B) distance to the nearest forest, and (C) forest area across the four forest dynamics categories. Each boxplot displays the median (central line), the interquartile range (IQR, the box), and the range of values (whiskers), with populations indicated as solid black dots.

To test whether population size can be used to infer extinction debt, population size was modeled against the log-transformed and scaled (mean-centered and divided by the standard deviation) forest area, scaled distance to the nearest forest and scaled latitude using GLMM with a negative binomial distribution in the *glmmTMB* package This model revealed no significant influence of forest dynamics or spatial variables (Table S5; Fig. S7).

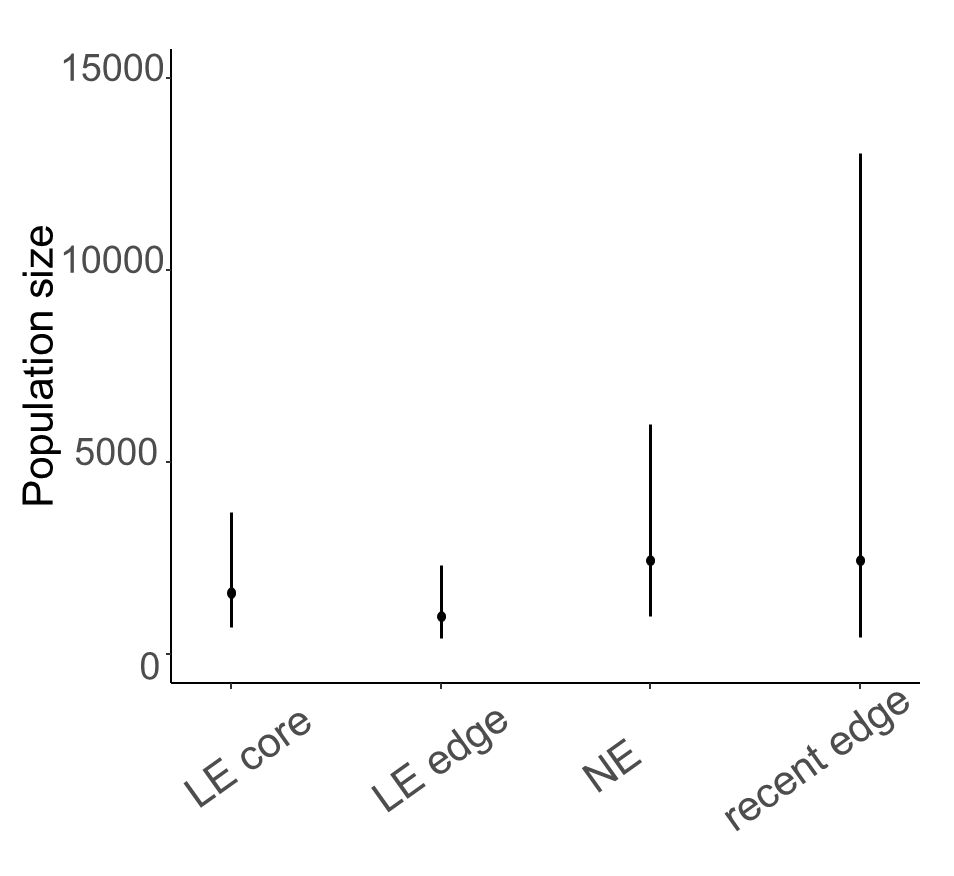

Figure S7: Prediction of the population size in relation to the forest dynamics based on the GLMM. Black dots show the average genetic diversity per individual factor, whereas error bars show the standard deviation. Abbreviations: LE = long-established, NE = newly established.
